# Different brain networks underlying intelligence in Autism Spectrum Disorders

**DOI:** 10.1101/143891

**Authors:** Emmanuel Peng Kiat Pua, Charles B Malpas, Stephen C Bowden, Marc L Seal

**Author notes:** Corresponding Author Emmanuel Pua Developmental Imaging, Murdoch Children’s Research Institute, The Royal Children’s Hospital, VIC 3052, Australia.

## Abstract

**Background:** There has been sustained clinical and cognitive neuroscience research interest in how network correlates of brain-behaviour relationships might be altered in Autism Spectrum Disorders (ASD) and other neurodevelopmental disorders. As previous work has mostly focused on adults, the nature of whole-brain connectivity networks underlying intelligence in pediatric cohorts with abnormal neurodevelopment requires further investigation.

**Methods:** We used network-based statistics (NBS) to examine the association between resting-state functional Magnetic Resonance Imaging (fMRI) connectivity and fluid intelligence ability in male children (*n*=50) with Autism Spectrum Disorders (ASD; *M*=10.45, *SD*=1.58 years and in controls (*M*=10.38, *SD*=0.96 years) matched on fluid intelligence performance, age and sex. Repeat analyses were performed in independent sites for validation and replication.

**Results:** Despite being equivalent on fluid intelligence ability to strictly matched neurotypical controls, boys with ASD displayed a subnetwork of significantly increased associations between functional connectivity and fluid intelligence. Between-group differences remained significant at higher edge thresholding, and results were validated in independent-site replication analyses in an equivalent age and sex-matched cohort with ASD. Regions consistently implicated in atypical connectivity correlates of fluid intelligence in ASD were the angular gyrus, posterior middle temporal gyrus, occipital and temporo-occipital regions.

**Conclusion:** Development of fluid intelligence neural correlates in young ASD males is aberrant, with an increased strength in intrinsic connectivity association during childhood. Alterations in whole-brain network correlates of fluid intelligence in ASD may be a compensatory mechanism that allows equal task performance to neurotypical peers.

## Introduction

Fluid intelligence refers to the ability to solve novel problems, and is typically estimated from composite scores of non-verbal or abstract tests. Under the Catell-Horn-Carroll (CHC) model, fluid intelligence (Gf) is a broad ability subsumed under an overall general intelligence factor (Reynolds and Keith, 2017; Schneider and McGrew, 2012). The autism spectrum disorders (ASD) are a group of heterogeneous neurodevelopmental conditions associated with deficits in social communication, social interaction, and restricted and repetitive behaviours. Uneven subtest scores within IQ measures have been observed across high- and low functioning ASD populations, as well as in both children and adults with ASD (Happé, 1994). In particular, non-verbal fluid intelligence performance in ASD groups has been suggested to be elevated relative to crystallized intelligence scores on verbal tasks. However, a neurobiological account of brain-behaviour associations in intelligence ability in ASD is lacking, and the link between altered neurodevelopment in ASD and cognitive performance remains poorly understood (Ehlers, et al., 1997; Happé, 1994; Hayashi, et al., 2008).

Previous investigations on the neural correlates of ASD fluid intelligence ability have primarily relied on task-based paradigms in functional Magnetic Resonance Imaging (fMRI), based on blood oxygen level dependent (BOLD) response as an estimate of local brain regions recruited when performing a particular task. Adults with ASD demonstrate an abnormal reliance on enhanced visuospatial processes in extrastriate and parietal regions when engaging in fluid tasks (Koshino, et al., 2005; Mottron, et al., 2013). Increase in fluid task complexity modulated stronger activity in occipital and temporal regions, coupled with higher connectivity between major lobar regions (Simard, et al., 2015; Soulières, et al., 2009). Fluid intelligence performance in neurotypical individuals is known to involve broad recruitment across frontal, parietal, temporal and occipital cortices, as well as subcortical striatal and thalamic regions (Burgaleta, et al., 2014; Geake and Hansen, 2010; Gong, et al., 2005; Kroger, et al., 2002; Perfetti, et al., 2009; Prabhakaran, et al., 1997). In contrast, connectivity to prefrontal cortical areas observed in controls during fluid tasks were either altered or absent in adults with ASD, suggesting impairments in functional segregation and integration in neural mechanisms underlying ASD fluid intelligence ability primarily characterized by increased occipito-parietal and temporal activity (Sahyoun, et al., 2010; Yamada, et al., 2012).

Previous studies focused on localization may be associated with a bias towards identification of task-positive regions, and there is a need for network-based whole-brain investigations (Basten, et al., 2015; Langeslag, et al., 2013). One approach to elucidate the neural correlates of fluid intelligence is the use of resting-state or intrinsic functional connectivity, defined as the temporal synchronicity of BOLD time-series between spatially distinct brain regions when no task is being performed (Lowe, et al., 2016). Intrinsic functional connectivity offers powerful methods to characterize the functional architecture of the brain. The measure corresponds to individual differences during task-dependent active states, and has also been shown to predict fluid intelligence ability (Smith, et al., 2009; Tavor, et al., 2016). However, most investigations on brain networks underlying cognition tend to be limited to general intelligence or neurotypical adult populations (Haász, et al., 2013; Malpas, et al., 2016; Penke, et al., 2012). The nature of whole-brain fluid intelligence connectivity networks in neurodevelopmental disorders is thus not well understood in paediatric populations. Specifically, it is not clear how the neurocognitive architecture of fluid intelligence is altered in ASD during early stages of cognitive development (Gray, et al., 2003; Kosslyn, et al., 2002).

To address this gap, we investigated whether whole-brain intrinsic functional connectivity networks associated with fluid intelligence were altered in children with ASD. Comparisons for all analyses were made to typically developing controls matched on age, sex and fluid intelligence ability. Importantly, strict matching criteria was implemented to ensure that observed differences were not likely to be explained by common confounds faced in ASD research (Pua, et al., 2017).

## Materials and Methods

### Participants

Data was obtained from the Kennedy Krieger Institute (KKI, ABIDE-II) sample from the Autism Brain Imaging Database Exchange (ABIDE; Di Martino, et al., 2014). ABIDE sites were only selected if comprehensive fluid intelligence measures were available, and age ranges and sample sizes were suitable for analyses (See Supplementary Figure 1). Following this criteria, the KKI site was included for the main analysis. Participants in the KKI sample were recruited as part of a study run by the Center for Neurodevelopment and Imaging Research (CNIR) at the KKI. All eligible participants received an MRI scan and cognitive assessment with the Wechsler Intelligence Scale for Children (*Fourth Edition,* WISC-IV; *Fifth Edition*, WISC-V). Handedness was assessed using the Edinburgh Handedness Inventory. Inclusion criteria were an age range of 8 years and 0 months to 12 years, 11 months and 30 days, and WISC-IV or WISC-V Full Scale Intelligence Quotient >80. Diagnosis of ASD was determined using the Autism Diagnostic Interview-Revised (ADI-R), Autism Diagnostic Observation Schedule-Generic (ADOS-G) module 3 or the ADOS-2 module 3. ASD classification criteria was based on the ADOS-G and/or ADI-R and clinical assessment by an expert pediatric neurologist with extensive experience in autism diagnosis. ASD participants were excluded if they had an identifiable cause of autism. For the control group, participants with a history of developmental or psychiatric disorders or with a first-degree relative with ASD were excluded. Full protocol details for sampling, image acquisition and phenotyping for all ABIDE sites are available elsewhere^1^.

For the present study, inclusion criteria applied to the KKI sample were male participants satisfying DSM-IV-TR^2^ Pervasive Developmental Disorder criteria (Autistic Disorder, Asperger’s or Not Otherwise Specified), assessed with the WISC-IV, and with MRI data acquired under the same scanning protocol. Continuous variables in the phenotype data were demeaned. Non-parametric propensity matching was performed using the MatchIt package (Ho, et al.) in the R environment (Team, 2014). Male participants with ASD were matched with TD controls on the following variables: sex, age in years [ASD: *M*=10.45 (*SD*=1.58); TD: *M*=10.38 (*SD*=0.96)], and PRI score (Perceptual Reasoning Index, or fluid intelligence score estimate) from the WISC-IV [ASD: 108.65 (13.57); TD: 108.83 (14.14)]. The matching procedure resulted in a final sample of 50 male participants (ASD: n=26; TD: n=24).

To evaluate if findings could be replicated, we repeated the same image processing and analysis on independent samples from other ABIDE sites. The Georgetown University (GU) site was selected based on similarity to the KKI sample in the main analyses in cohort age range (8 to 13 years), availability of fluid intelligence measures, and adequate sample size. Because imaging acquisition protocols and parameters, and behavioural test measures differed between sites, GU was used as a validation sample rather than collapsed with the initial analyses. To investigate if similar results could be observed in an older age cohort (age range: 15 to 24 years), the same analysis was repeated in The University of Utah School of Medicine (USM) sample. Table 1 provides descriptive statistics for all samples used for present analyses.

**Table 1.** Descriptive statistics of samples by site

| Site | Sample size | Age in years [M(SD)] | Age range in years | PRI [M(SD)] | Test type |
| --- | --- | --- | --- | --- | --- |
| **KKI** |  |  |  |  |  |
| ASD | 26 | 10.45 (1.58) | 8 to 13 | 108.65 (13.57) | WISC-IV |
| TD | 24 | 10.38 (0.96) | 8.9 to 12.8 | 108.83 (14.14) | WISC-IV |
| **GU** |  |  |  |  |  |
| ASD | 25 | 11.04(1.44) | 8.28 to 13.08 | 109.68 (14.06) | WASI |
| TD | 25 | 10.6(1.46) | 8.06 to 12.7 | 118.52 (13.45) | WASI |
| **USM** |  |  |  |  |  |
| ASD | 15 | 20.32 (1.48) | 18.41-22.88 | 102.67 (13.98) | WASI |
| TD | 15 | 18.46 (2.45) | 15-23.95 | 111 (10.39) | WASI |

*Notes.* ASD: Autism Spectrum Disorders; GU: Georgetown University; KKI: Kennedy Krieger Institute; M= mean; PRI: Perceptual Reasoning Index, or fluid intelligence score estimate; SD=standard deviation, TD: typically developing; USM: The University of Utah School of Medicine; WASI: Wechsler Abbreviated Scale of Intelligence; WISC-IV: Wechsler Intelligence Scale for Children – Fourth Edition.

### Image Acquisition

MRI data was acquired on a 3 Tesla Phillips scanner (Achieva; Philips Healthcare, Best, The Netherlands). T1-weighted images were obtained through a 200 slice three-dimensional acquisition (Turbo Field Echo [TFE] MPRAGE; acquisition time = 8min 8sec; coronal slice orientation; flip angle =8°; repetition time (TR) = 8ms; echo time (TE) = 3.7ms; minimum inverse time (TI) delay = 843.25ms; field of view (FOV) = 256; matrix= 256 x 200; slice thickness = 1mm, in-plane resolution = 1mm x1.28mm). BOLD-weighted resting-state functional MRI volumes were acquired using echo planar imaging (EPI; number of volumes = 128; TE = 30ms; TR = 2500ms; slices = 47; flip angle = 7°; FOV = 256; matrix = 84×81; slice thickness = 3mm, in-plane resolution = 3mm x3.1mm; transverse slice orientation). During the resting-state scan, participants were instructed to relax and focus on a crosshair while remaining as still as possible with their eyes open. For the structural scan, participants watched a movie of their choice. Images were inspected after each processing step for quality control.

### Image Processing and Analysis

Functional connectivity analysis and visualizations were generated with the Functional Connectivity Toolbox v.16.b (Whitfield-Gabrieli and Nieto-Castanon, 2012) pipeline, Matlab R2010b (The MathWorks, Inc., Natick, MA, USA) and NeuroMArVL (http://immersive.erc.monash.edu.au/neuromarvl/). The initial 4 functional volumes per session were removed to account for T1 saturation effects. Slice-timing correction and first-volume realignment (using a six rigid-body parameter spatial transformation) were applied to adjust for temporal and motion artefacts. Functional volumes were normalized to MNI-space, and smoothed with a full-width half-maximum Gaussian kernel of 8mm. Structural images were co-registered and segmented into grey-matter, white-matter and cerebrospinal fluid for later use in the removal of physiological noise from the functional volumes. In the first-level BOLD model, realignment transformation matrices and global signal intensities were analyzed using the Artifact Detection Tool (ART) to detect motion and signal outliers. Noise from sources such as cardiac, respiratory, and other physiological activity that were not likely to be modulated by neural activity were identified using the component-based noise correction aCompCor method (Behzadi, et al., 2007). Confounds were removed by regressing artefactual effects from the BOLD signal (within-subject covariates from realignment parameters, non-neuronal physiological noise from white-matter and cerebrospinal fluid masks). Residual BOLD time series were detrended and band-pass filtered (0.008-0.09Hz) before computing connectivity measures. Regions-of-interest (ROI) were defined using the FSL Harvard-Oxford Atlas (http://www.fmrib.ox.ac.uk/fsl/) for cortical and subcortical areas, and the Anatomical Automatic Labelling (AAL) atlas (Tzourio-Mazoyer, et al., 2002) for cerebellar regions, resulting in 132 ROIs. The mean BOLD time series for all voxels in each ROI were extracted to compute pairwise correlations between all ROIs with the Fisher r-to-z transformation to construct a 132×132 connectivity matrix.

Network-based statistic (NBS) was used to identify brain networks showing between-group differences in functional connectivity associated with fluid intelligence performance (Zalesky, et al., 2010). NBS is a network-specific approach that is typically used to identify groups of connections showing a significant effect of interest, while controlling for the family-wise error (FWE) rate. As a nonparametric method for connectome-wide analysis, NBS offers substantially greater statistical power than generic mass-univariate testing procedures at the edge level, if connections associated with the effect of interest form connected components (defined here as interconnected subnetworks). NBS estimates an FWE-corrected *p*-value for identified subnetworks through permutation testing. Consequently, the null hypothesis can only be rejected at the level of entire subnetworks, and inferences about specific connections will not be valid. The principle behind NBS is similar to cluster-based approaches used to perform inference on statistical parametric maps, and utilizes the observation that pathological effects of disease are frequently distributed across the brain but also constrained by anatomical network topology (Fornito, et al., 2016). NBS has been widely used to study altered brain connectivity in disorders such as schizophrenia (Zalesky, et al., 2011), depression (Korgaonkar, et al., 2014), addiction (Hong, et al., 2013), Parkinson’s disease (Aarabi, et al., 2015), as well as connectivity correlates of cognition in the general population (e.g. Hearne, et al., 2016b).

For the present study, fluid intelligence performance scores were first regressed onto individual edges in the functional connectivity matrix. Between-group differences in functional connectivity associated with performance scores were evaluated at the second-level. Handedness and age were included as covariates for all analyses. To identify interconnected subnetworks across groups, a breadth first search (Ahuja, et al., 1993) was performed among connections surviving a t-statistic threshold (i.e the component forming threshold) of at least t=3.0, and permuted to generate a null distribution of largest network sizes. Each permutation randomly reassigns group labels and identifies the size of the largest interconnected subnetwork to yield an empirical null distribution of maximal component sizes. The estimated FWE-corrected p-value for an identified subnetwork of size *m* reflects the proportion of permutations for which the largest subnetwork size is equal to or greater than *m*. The FWE rate is therefore controlled nonparametrically using a randomized null distribution of maximum component size. Subnetworks with a corrected p-FWE<0.05 value were retained.

## Results

Between-group differences in the association of resting-state fMRI subnetwork connectivity with fluid intelligence performance was identified in males aged 8 to 13 years (network size=24 links, *p*=.0373, FWE-corrected), with stronger association with fluid intelligence in ASD compared to typically developing matched controls (Figure 1). This network showing stronger correlation with fluid intelligence in ASD compared to TD involved regions in the left temporo-occipital middle temporal gyrus, left posterior middle temporal gyrus, bilateral paracingulate gryus, posterior cingulate gyrus, right frontal pole, right inferior frontal gyrus pars triangularis, bilateral angular gyrus, left lateral occipital cortex (superior division) and precuneus (Table 2). Between-group differences remained significant even at a more stringent component-defining threshold (t=|4|, size=6, p=.0425, FWE-corrected), with nodes of the left temporo-occipital middle gyrus, bilateral angular gyrus, precuneus, posterior cingulate gyrus that survived more conservative analyses.

**Figure 1.**
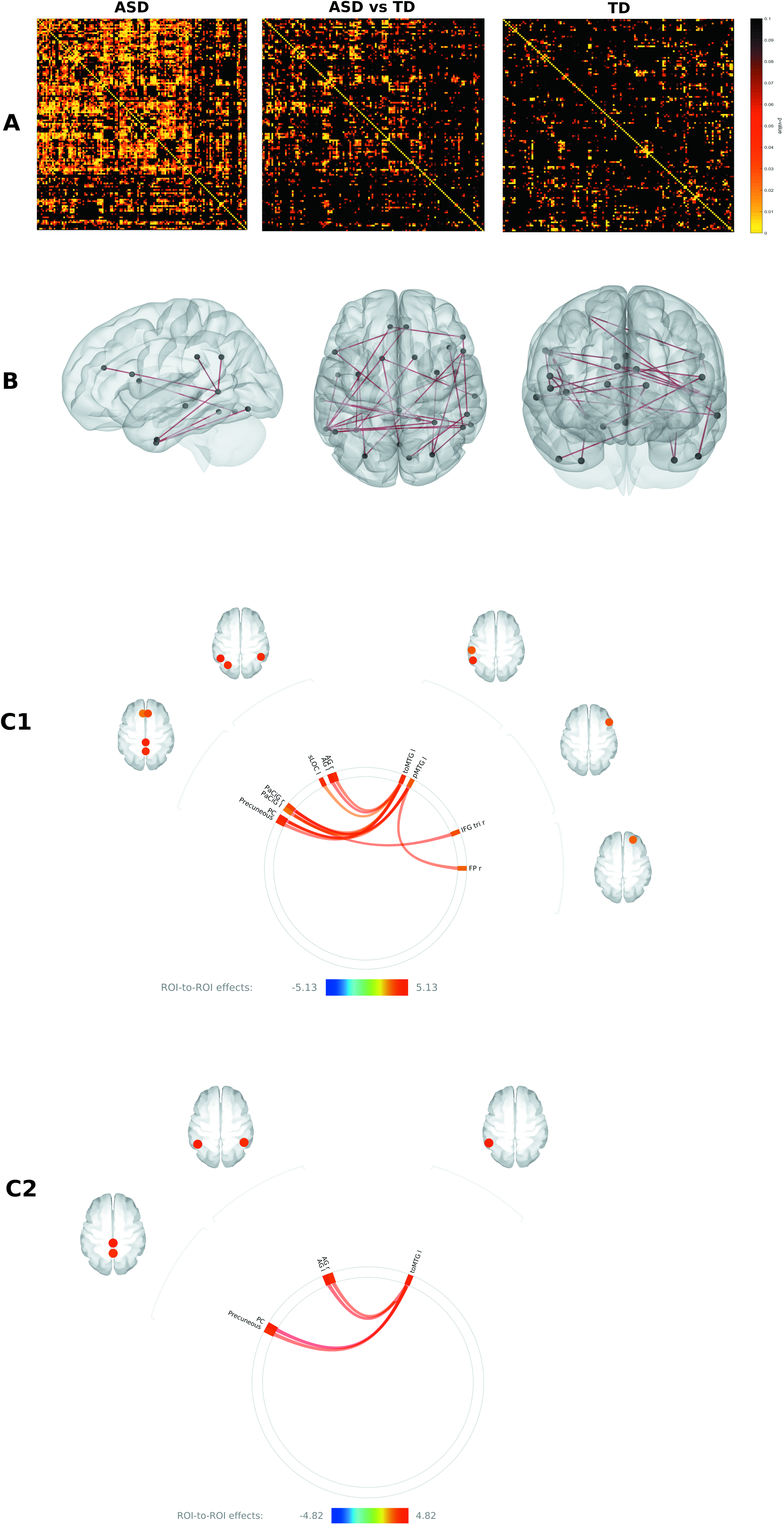
**A**: P-value heatmaps representing second-level within-group associations of pairwise BOLD ROI-ROI edges (132×132 matrix) with fluid intelligence performance (Site: KKI). ASD: Autism Spectrum Disorders group; ASD vs TD: Group interactions between functional connectivity and fluid intelligence performance association; KKI: Kennedy Krieger Institute; TD: matched typically developing controls. **B:** Visual model of subnetwork representing within-group associations in functional connectivity and fluid intelligence performance in ASD. **C:** Effect of increased thresholding in network-based analysis (NBS) before network construction. In C1, only pairwise edge association with fluid intelligence of at least t=3.5 were retained for network construction. In C2, the t-statistic threshold was further increased to t=4. AG: angular gryus; FP: frontal pole; IFG tri: inferior frontal gyrus, pars triangularis; PaCiG: paracingulate gyrus; PC: posterior cingulate gyrus; sLOC: Lateral occipital cortex, superior division; toMTG: temporo-occipital middle temporal gyrus.

**Table 2.** Nodes identified in atypical subnetwork connectivity association with fluid intelligence in Autism Spectrum Disorders compared to matched controls (Site KKI)

| Lobe | Node | Position | Node Label | MNI Coordinates |  |  |
| --- | --- | --- | --- | --- | --- | --- |
|  |  |  |  | x | y | z |
| Frontal | Paracingulate gryus | Left | PaCiG l | -6.21 | 36.65 | 20.79 |
|  | Paracingulate gryus | Right | PaCiG r | 6.55 | 36.57 | 22.69 |
|  | Frontal pole | Right | FP r | 26.16 | 52.14 | 8.26 |
|  | Inferior frontal gyrus, pars triangularis | Right | IFG tri r | 51.87 | 27.76 | 7.71 |
| Temporal | Middle temporal gyrus, posterior division | Left | pMTG l | -60.91 | -27.36 | -11.00 |
|  | Temporo-occipital middle temporal gryus* | Left | toMTG l | -57.64 | -53.00 | 0.82 |
| Occipital | Lateral occipital cortex, superior division | Left | sLOC l | -31.96 | -72.89 | 37.97 |
| Parietal | Angular gyrus* | Right | AG r | 51.93 | -51.80 | 32.36 |
|  | Angular gyrus* | Left | AG l | -50.35 | -55.70 | 29.76 |
|  | Precuneous* | - | Precuneous | 0.95 | -59.29 | 38.02 |
|  | Posterior cingulate gyrus | - | PC | 0.78 | -36.62 | 29.98 |

Notes. \*Nodes surviving increased thresholding (T=4) in network-based statistics analysis (NBS); KKI: Kennedy Krieger Institute.

Repeat analyses in samples from independent sites presented a similar pattern of findings in a cohort of comparable age (site GU; age range 8 to 13 years), showing a single fluid intelligence subnetwork with increased association in ASD compared to controls (network size=14 links, t-statistic threshold=3.5, *p*=.0396, FWE-corrected, alternate hypothesis: ASD>controls; Figure 2). Implicated regions were the bilateral occipital pole, right temoporo-occipital middle temporal gyrus, right anterior middle temporal gyrus, left posterior middle temporal gyrus, right angular gyrus and the cerebellum (Table 3). Across all analyses from independent sites in age-matched samples, the right angular gyrus, left posterior middle temporal gyrus, occipital and temporo-occipital regions were consistently implicated in fluid intelligence subnetwork differences. No subnetworks were identified under the alternate hypothesis in the opposite direction (controls > ASD). In contrast to findings from the younger age cohorts, no subnetworks of increased connectivity associated with fluid intelligence were identified in older individuals with ASD ages 15 to 24 years (site USM; p>.05, FWE-corrected). No subnetworks were identified even when the initial thresholding of pairwise functional connectivity links and FWE-corrected p-values were relaxed.

**Table 3:** Nodes identified in atypical subnetwork connectivity association with fluid intelligence in Autism <u>Spectrum Disorders compared to matched controls (Site GU)</u>

| Lobe | Node | Position | Node Label | MNI Coordinates |  |  |
| --- | --- | --- | --- | --- | --- | --- |
|  |  |  |  | x | y | z |
| Frontal | - | - | - | - | - | - |
| Temporal | Middle temporal gyrus, anterior division | Right | aMTG r | 57.89 | -1.529 | -24.51 |
|  | Middle temporal gyrus, posterior division* | Left | pMTG l | -60.91 | -27.36 | -11.00 |
|  | Temporo-occipital middle temporal gyrus* | Right | toMTG r | 58.18 | -49.22 | 1.59 |
| Occipital | Occipital pole | Right | OP r | 17.73 | -95.13 | 8.31 |
|  | Occipital pole | Left | OP l | -16.85 | -96.50 | 6.74 |
| Parietal | Angular gyrus* | Right | AG r | 51.93 | -51.80 | 32.36 |
| Cerebellar | Cerebellum, crus II | Left | Cereb2 l | -28.64 | -73.26 | -38.20 |
|  | Cerebellum, vermis 8 | - | Ver8 | 1.15 | -64.43 | -34.08 |

Notes. \*Nodes consistent across independent-site replication GU: Georgetown University

**Figure 2.**
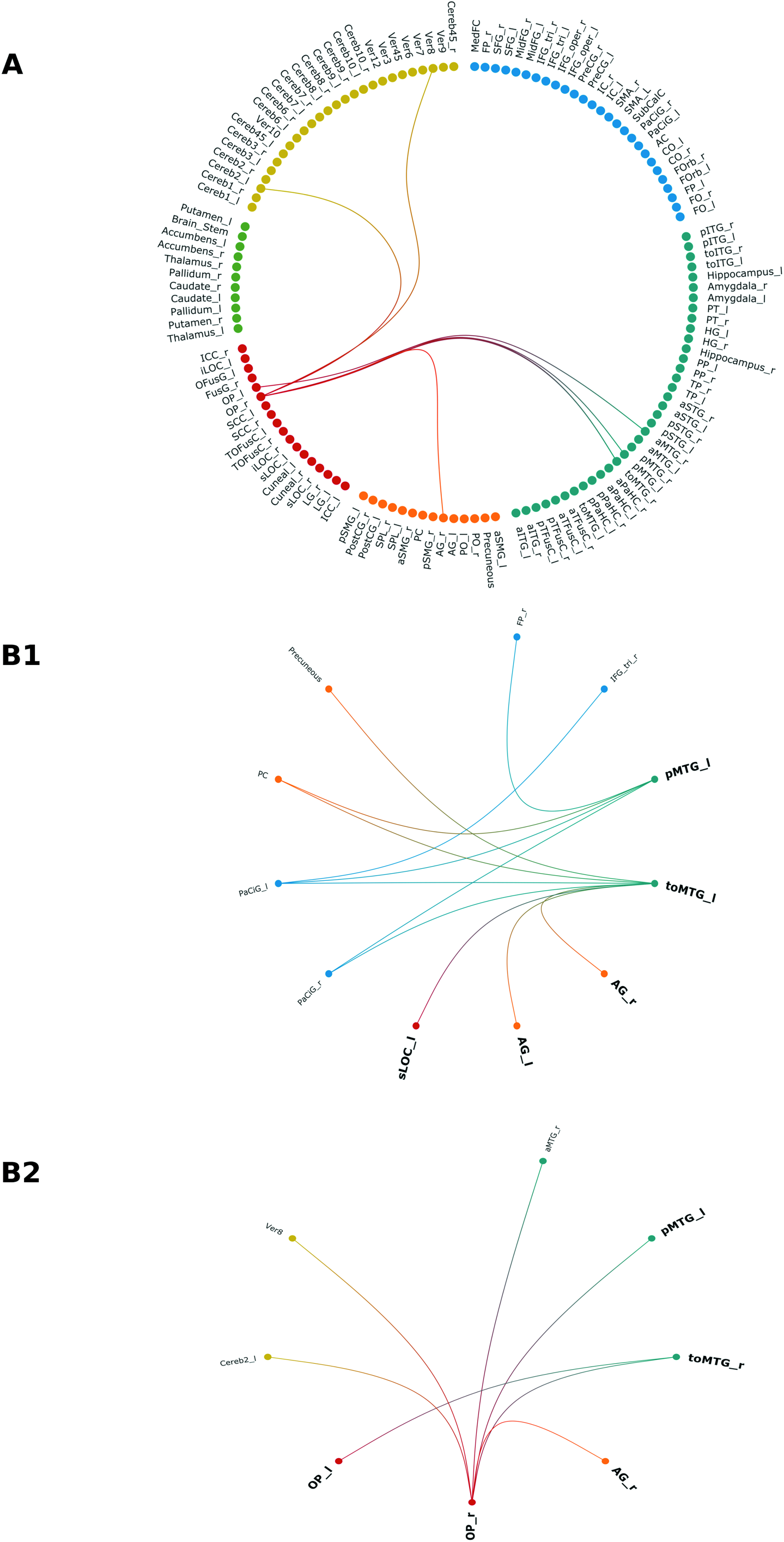
**A**: Cortical and subcortical regions involved in atypical neural correlates of ASD fluid intelligence in replicated analyses, configured by lobe. The subnetwork represents between-group differences in edge connectivity and fluid intelligence performance association in ASD compared to matched controls (Site: Georgetown University, GU). **B:** Network components implicated in atypical neural correlates of ASD fluid intelligence (B1; Site: Kennedy Krieger Institute, KKI) and in independent-site replication (B2; Site GU). AG: angular gryus; aMTG: anterior middle temporal gyrus; Cereb2: cerebellum, crus II; FP: frontal pole; IFG: inferior frontal gyrus; OP: occipital pole; PaCiG: paracingulate gyrus; PC: posterior cingulate gyrus; pMTG: posterior middle temporal gyrus; sLOC: Lateral occipital cortex, superior division; toMTG: temporo-occipital middle temporal gyrus; Ver8: cerebellar vermis 8.

## Discussion

This is the first study to investigate the neural correlates of fluid intelligence in paediatric ASD cohorts based on a novel brain-behaviour matching framework. We identified increased resting state functional connectivity associated with fluid intelligence ability in young ASD males relative to matched controls. These results were replicated in an independent sample with a matched cohort of the same age range. In the context of inter-site differences in sampling, phenotyping, diagnostic classification, MRI scanner and acquisition parameters, and significant heterogeneity within ASD neurobiology, the robustness of present findings suggest the presence of aberrant functional connectivity correlates of fluid intelligence in ASD between 8 to 13 years of age.

Brain regions identified are consistent with previous investigations of localized functional BOLD activity during fluid task performance that show atypical increases in occipito-temporal activity coupled with decreased prefrontal activation in ASD (Yamada, et al., 2012). A recent meta-analysis of grey-matter abnormalities in paediatric ASD reported grey-matter alterations in the right angular gyrus, left inferior occipital gyrus and right inferior temporal gyrus, as well as in frontal, medial parietal and cerebellar regions. Increased grey-matter volume of the right angular gyrus was further associated with increased severity of repetitive behaviours, a core symptom in ASD (Liu, et al., 2017).

In neurotypical individuals, independent component analysis of task-based fMRI metadata show that temporo-occipital and inferior parietal regions constitute a cluster of intrinsic connectivity networks related to visual perception of complex stimuli higher-level visual processing, visual tracking, mental rotation and spatial discrimination (Laird, et al., 2011). The angular gyrus and temporal-occipital cortex also demonstrate shared activity even among different fluid intelligence tasks, suggesting their role as potential neural correlates of fluid intelligence ability (Ebisch, et al., 2012). In task-free states, connectivity between the angular gyrus and occipital regions form part of a temporally independent functional mode (Smith, et al., 2012). Functionally, the angular gyrus serves as a cross-modal hub that combines and integrates multisensory information for attentional reorientation to critical information, comprehension of environmental events, manipulation of mental representations and problem solving (Seghier, 2013). According to the P-FIT (Parieto-Frontal Integration Theory) hypothesis, the role of the angular gyrus in information processing stages involves integration and abstraction, followed by parietal-frontal interactions that support problem solving and evaluation of solutions (Colom, et al., 2009; Jung and Haier, 2007). That our findings cohere with known functional connectivity networks of fluid ability in typical controls suggest that these systems also subserve similar functions in ASD, but are more susceptible to aberrations in local and global functional connectivity due to atypical neurodevelopment.

Abnormalities in intrinsic connectivity associated with fluid intelligence were found in younger but not older age groups in the present study. Alterations in brain structure and function in ASD tend to be age-dependent, and are characteristic of atypical neurodevelopment in the disorder (Uddin, et al., 2013). In children with ASD, there is significant hypoactivation of the middle frontal gyrus during nonsocial tasks in contrast to adults with ASD (Dickstein, et al., 2013). A similar pattern of early increased functional connectivity followed by a decline in later stages has been reported in other neurodevelopmental disorders, possibly related to dysregulation of brain activity due to aberrant neurodevelopment of structural connectivity of hub regions in the association cortices (Fornito and Bullmore, 2015). As expected, the inconsistency of abnormalities across different ages presents a significant source of with within- and between-group heterogeneity that is difficult to capture reliably.

## Implications

It is of interest that between-group differences in intrinsic connectivity networks associated with fluid intelligence performance were observed between ASD and controls, despite both groups being equivalent on fluid intelligence ability. The degree of association between subnetwork connectivity and fluid ability observed in ASD is therefore more likely related to disorder-specific effects, rather than differences in fluid performance ability (Gray, et al., 2003; Perfetti, et al., 2009). According to the neural efficiency hypothesis, differential cortical activity can be observed among subjects with discrepant neural resources, despite similar task ability. Abnormally increased activation of brain regions during cognitive tasks in atypical populations may indicate a mechanism of neural compensation to achieve the same degree of performance. Consistent with present findings, the process often involves mediation by hub nodes that integrate multiple neural systems, such as the angular gryus in the parietal association cortex. Dedifferentiation, the failure of neural processes to specialize due to neurodevelopmental abnormalities could also underlie early aberrant increases in hub activity observed in our findings (Fornito, et al., 2017). Under this framework of neural compensation, our results also complement previous findings of increased BOLD signal changes with increased fluid task difficulty in the inferior parietal lobule including the angular gyrus, and the left temporo-occipital junction in healthy individuals (Preusse, et al., 2011). Increased resting-state connectivity was also associated with higher intelligence scores (Hearne, et al., 2016a). Together, atypically increased strength of association in the ASD fluid intelligence subnetwork may reflect a compensatory effect to achieve the same level of fluid task performance as ability-matched controls in our analyses.

Consequently, the common assumption that matching on group variables is sufficient to control for all associated variation is problematic, and could introduce artefactual differences in case-control comparisons biased by differential associations in neural correlates (Lefebvre, et al., 2015). As we have shown in this study, group differences in brain structure and function should also demonstrate covariation with variables of interest such as clinical symptom severity. Understanding how alterations in the brain relate to individual differences in phenotypic expression in ASD may reveal important pathways in the aetiology of ASD (Picci, et al., 2016). These cohort-dependent findings in ASD highlight the importance of ensuring that differences observed are associated with the effect of interest, rather than variation related to confounds (Bölte, et al., 2009; Dawson, et al., 2007). Given that we have investigated higher functioning ASD subgroups, further work should determine if similar patterns of alterations can be observed in low functioning individuals.

A final consideration are the constraints of NBS. The technique yields increased power over link-based FWE control to detect connected components with whole-brain multiple comparisons, but at the cost of localizing resolution for independent links (Zalesky, et al., 2010). We have thus refrained from directly interpreting individual edge links in identified subnetworks. Connectivity strength and network topology are distinct properties of the brain connectome that can demonstrate mutually exclusive perturbations (Hong, et al., 2013). While we have focused on functional connectivity analyses, our findings establish a promising framework for subsequent investigations into the multi-scale configuration of the neural correlates of cognition in various disorders, and across different neuroimaging modalities. As our pipeline only included one method of parcellation, replication of analyses with different parcellation techniques could also strengthen present findings by reducing the likelihood of atlas selection biases (Abraham, et al., 2016).

Apart from identifying fundamental units or collective features of network topology, graph-based measures allow the identification of intermediate mesoscale structures through community detection techniques and across multiple timescales. Importantly, the function of network nodes may differ depending on the scale of analysis (Betzel and Bassett, 2016). Given also that structural connectivity correlates were not investigated in the present study, cross-modal network analyses that integrate structural and functional data will be crucial to delineate the mechanisms of cognitive development, and how the nature and developmental trajectories of the neural correlates of cognition are altered over time in neurodevelopmental conditions (Grayson and Fair, 2017). Because anatomical networks determine pathways of neuronal signaling, structural network development likely precedes the complete deployment of global intrinsic functional connectivity networks underlying intellectual ability in children and adolescents. (Petersen and Sporns, 2015; Vertes and Bullmore, 2015). The characterization of the neural architecture of intelligence in both typical and atypical neurodevelopmental populations requires further exploration of brain structural and functional network connectivity and topology across multiple scales, and the integration of neuroimaging findings with valid measurement of cognitive constructs of interest.

## Conclusion

We demonstrate novel preliminary evidence with replication for an atypical intrinsic connectivity subnetwork associated with fluid intelligence in male ASD children compared to controls. For this particular age cohort, the neural architecture of fluid intelligence in ASD children involves aberrant network integration of distributed regions and an increased strength of association to support equal task performance with same-aged peers. There is potential for longitudinal investigations to delineate inter- and intra-individual variation and between-sex differences in the neurodevelopment of cognitive ability across different populations.

## Acknowledgments

Data analysis and interpretation was conducted within the Developmental Imaging research group, Murdoch Children’s Research Institute and the Children’s MRI Centre, Royal Children’s Hospital, Melbourne, Victoria. All procedures were conducted in accordance with the Declaration of Helsinki, and approved by The Royal Children’s Hospital Human Research Ethics Committee. Data used in this research study was acquired from the public-access Autism Brain Imaging Data Exchange (ABIDE; http://fcon_1000.projects.nitrc.org/indi/abide/) funded by the National Institute of Mental Health (NIMH 5R21MH107045), and available through the link provided.

The research was supported by the Murdoch Children’s Research Institute, the Royal Children’s Hospital, Department of Paediatrics The University of Melbourne and the Victorian Government’s Operational Infrastructure Support Program. The project was generously supported by RCH1000, a unique arm of The Royal Children’s Hospital Foundation devoted to raising funds for research at The Royal Children’s Hospital. The authors declare no potential conflicts of interest with respect to the research, authorship, and/or publication of this article.

## Footnotes

1 Public access ABIDE protocol: http://fcon_1000.projects.nitrc.org/indi/abide/

2 DSM-IV-TR: Diagnostic and statistical manual of mental disorders, text revision. American Psychiatric Association, & American Psychiatric Association. (2000). DSM-IV-TR: Diagnostic and statistical manual of mental disorders, text revision. Washington, DC: American Psychiatric Association, 75.

